# Genetic instability as a driver for immune surveillance

**DOI:** 10.1101/527689

**Authors:** Guim Aguadé-Gorgorió, Ricard Solé

**Affiliations:** ICREA-Complex Systems Lab, Universitat Pompeu Fabra, 08003 Barcelona, Spain and Institut de Biologia Evolutiva (CSIC-UPF), Psg Maritim Barceloneta, 37, 08003 Barcelona, Spain; ICREA-Complex Systems Lab, Universitat Pompeu Fabra, 08003 Barcelona, Spain Institut de Biologia Evolutiva (CSIC-UPF), Psg Maritim Barceloneta, 37, 08003 Barcelona, Spain and Santa Fe Institute, 399 Hyde Park Road, Santa Fe NM 87501, USA

## Abstract

Genetic instability is known to relate with carcinogenesis by providing tumors with a mechanism for fast adaptation. However, mounting evidence also indicates causal relation between genetic instability and improved cancer prognosis resulting from efficient immune response. Highly unstable tumors seem to accumulate mutational burdens that result in dynamical landscapes of neoantigen production, eventually inducing acute immune recognition. How are tumor instability and enhanced immune response related? An important step towards future developments involving combined therapies would benefit from unraveling this connection. In this paper we present a minimal mathematical model to describe the ecological interactions that couple tumor adaptation and immune recognition while making use of available clinical estimates of relevant parameters. The possible evolutionary trade-offs associated to both cancer replication and T cell response are analysed, indicating that cancer-clearance states become attainable when both mutational load and immune migration are enhanced. Furthermore, the model predicts the presence of well-defined transitions towards tumor control and eradication after increases in genetic instability consistent with available data of tumor control after Mismatch Repair knockout.

## I. Introduction

Cancer is a disease of Darwinian cellular evolution [1]. Following depletion of a vast set of genetic insults altering normal multicellularity phenotypes, rogue cells are able to adapt and evade selection barriers leading to uncontrolled proliferation. In this context, genomic instability plays a key role as a driver of the genetic novelties required for tumor progression and rapidly adapting phenotypes [2, 3]. High levels of evolving instability sustain a very diverse population [4], and intra-tumor heterogeneity lies at the very core of why cancer is still difficult to define, characterize and cure [5].

In this paper we aim at understanding an important relationship between the effectiveness of cancer immunotherapy and genetic instability. The relevance of such link needs to be found in the challenges faced by immunotherapies based on immune checkpoint inhibition or adoptive cell transfer [6], where mutational burden seems to play a key role. Due to the underlying complexity of cancer immunology, interdisciplinary efforts towards novel immunotherapies are much required [7]. As discussed below, the crucible of the problem might be associated to the nonlinear dynamics associated to cancer neoantigen production and the consequent enhancement of immune surveillance.

A key point in cancer immunotherapy lies on the mechanisms by which T cells actually recognize cancerous from healthy tissue [8] and eventually attack tumor cells expressing tumor-specific antigens [9]. On a general basis, such antigens can be common proteins for which T cell acceptance is incomplete, or more importantly, novel peptides absent for the normal human genome [8,10]. Except for specific tumor types of viral etiology, this so-called *neoantigens* arise after DNA damage resulting in the production of novel proteins. Recent advances highlight the importance of understanding neoantigen generation as a consequence of the tumor mutational load and dissecting specific neoantigen immunogeneicity [8,9,11]. Furthermore, direct correlations have been suggested between neoantigen production at high microsatellite instability and eventual immune surveillance and clinical response to immunotherapies [12–14].

Several experimental and clinical sources are pointing towards a causal relation, including tumor growth impairment after inactivation of MLH1 [15], or the positive response to PD-1 blockade across different mismatch repair (MMR) defficient cancer types [16]. The inactivation of MMR results in increased mutational burden of cancer cells, promoting the generation of neoantigens which improve immune surveillance and eventual tumor arrest. These perspectives suggest a novel view on immunotherapy, where targeting mutagenic pathways can result in an alternative mechanism to unleash immune responses.

All in all, genetic instability seems to play a conflictive role in cancer evolution and proliferation. It appears that the same lessions that activate cancer progression can on the other hand account for T cell recognition and immune attack. The extent of such trade-off and its application to therapy, however, is not clear. On the one hand, mutagenic therapies coexist with an intrisic risk, as increased genetic instability on heterogeneous populations might activate oncogenic outgrowth in previously stable cells. Together with this, a reactive immune system might pose a selective pressure for immune editing, translating as selection for T cell evading tumor subclones. How do these two components-instability and immune response- interact and what are the consequences? Is it possible to provide useful insight from mathematical models without a detailed picture of the immune landscape of cancer [17]?.

Nonlinear responses associated to cancer-immune system interactions have been known from the early days of cancer modelling, from more classical approaches [18] to recent perspectives based on neoantigen recognition fitness [19]. These studies have revealed a number of interesting properties exhibited by toy models, including in particular the existence of shifts and breakpoints separating cancer progression from its extinction (see [20] and references therein). Such shifts are of exceptional importance in our context: they indicate the existence of well defined conditions (and perhaps therapeutic strategies) allowing an all-or-none response. However, a mathematical description of the specific role of genetic instability in cancer immunology has not yet been developed. Below we provide a first approach to such goal, based on considering both cancer adaptation and immune surveillance as influenced by mutational burden, and we analyze how genetic instability can account for transitions towards states of cancer control and elimination. The implications of these transitions on both mutagenic and immunotherapies are discussed, pointing towards possible cross-therapies activating neoantigen production and immune stimulation.

## II. QUICK GUIDE TO MODEL AND MAJOR ASSUMPTIONS

### A. Mean-field cancer-immune model

The cellular interactions considered here involve a well-mixed (mean-field) model [20,21] where the population of cancer cells *c* follows a logistic growth (at replication rate *r* and carrying capacity *K*) and immune-cell mediated death (at rate *δ_c_*)

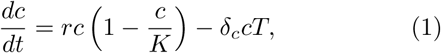

while the effector T cell dynamics are described by

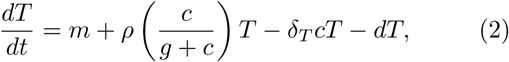

indicating a population that grows due to a constant migration of cells *m* and a predation term *ρ*, related with antigen recognition, that saturates at high tumor concentrations (*g*) due to limitations such as in the rate of T cell penetration. T cells have a natural decay rate, *d*, and also die when competing with tumor cells at rate –*δ_T_c*. The complete set of interactions described by (1) and (2) is shown schematically in figure 1.

### B. A one-dimensional landscape for tumor instability

Genetic instability has a twofold impact on cell fitness. As discussed along the introduction, replication rate *r* is a function of mutation probability *µ*. A landscape *r*(*µ*) is now in place [22,23], and follows from considering that mutations on oncogenes can translate into a linear increase in replication rate. This will be expressed as *R*_1_(*µ*) = *r*_0_ + *N_R_δ_R_µ* with *r*_0_ being the basal replication rate of normal cells, *N_R_* the number of oncogenes responsible for increased replication and *δ_R_* the mean effect on replication rate when mutating one of such genes.

**FIG. 1:**
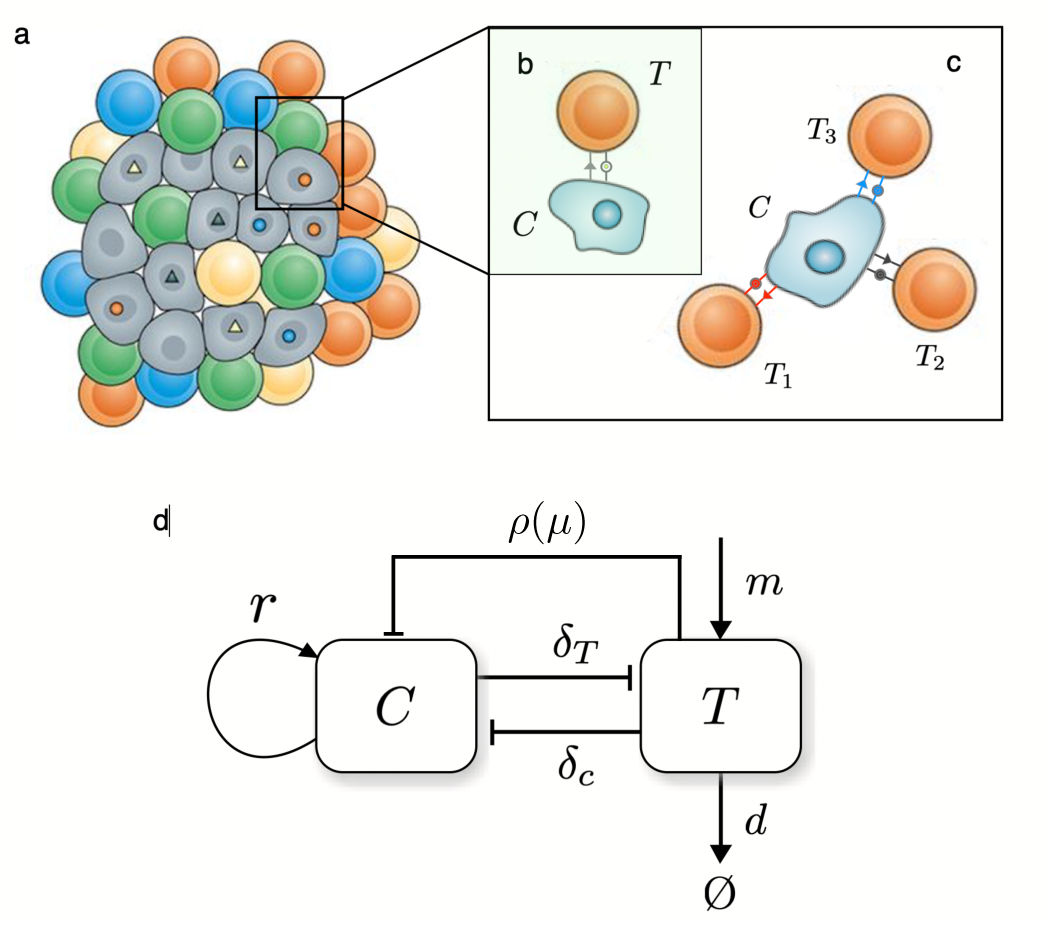
A schematic summary of the basic cancer-immune cell-cell interactions. The two key components are (a) a tumor population driven by genetic instability and (b-c) interactions associated to tumor cell recognition by T cells. The strength T cell attack depends on the number of surface neoantigens (b,c). In (d) the population-level interaction diagram is displayed based on the model in [20]. Here *C* and *T* indicate the number of cancer and T cells, respectively. Cancer cells grow at a rate *r* (and have a limited carrying capacity) while T cells enter the system at a constant production rate *m* and detect malignant cells at an instability-dependent rate *ρ*(*µ*). A constant death rate *d* is associated with their removal. Two constant cross-interactions rates are also indicated as *δ_T_* and *δ_c_* associated to the removal efficiency of cancer cells and the death of T cells resulting from the same process, respectively.

The number of house-keeping genes *N_HK_* is taken into account, and cell viability is conserved as long as none of them is mutated, with probability *R*_2_(*µ*) = (1 – *µ*)^*N_HK_*^. Grouping both considerations together we obtain an analytical description of the coupling between replication rate and mutation probability *r*(*µ*) = *R*_1_(*µ*)*R*_2_(*µ*) which reads:

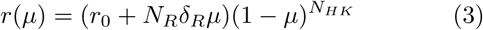

On the other hand, a linear approximation for malignant cell recognition by the immune system as a function of the tumor’s mutational load *µt* is obtained from previous data across many tumor types [24]: *ρ*(*µ*) = 4.35 · 10^−4^*µt*. However, we expect a function depending only on mutation probability. The variable *t* in this expression refers to the evolutionary life history of mutations accumulation of the tumor. This time scale is much larger than the faster ecological dynamics that govern the cancer-immune system interactions, so that we can consider it an average measure of tumor age at the time of detection, and consider it constant when introducing *ρ* in the ecological dynamics. From these facts, the only variable governing immune recognition at the cancer-immune competition level is the point mutation probability *µ*.

A very rough estimate for *t* can be either inferred from average cell replication data and from the fact that values for the mean mutation rate and the absolute mutational load are known for many tumors [25]. Taking a mean value of 0.18 divisions per day [20], an average tumor life in the range of about 10^1^ ~ 3 · 10^1^ and the gene number to be in the range of 2 · 10^4^, we obtain *t* ~ 10^7^ gene divisions per tumor life. This fits with the notion that mutator tumors might have mutation rates of about 10^−5^ mutations per gene division, which account for the accumulation of about 10^3^ somatic mutations per tumor life [26], so that tumor average tumor divisions lies again at about *t* ~ 10^7^. Using this approximation stemming from both methods we obtain our preliminar expression for how the immune reactivity rate depends on the mutation levels, *ρ*(*µ*) = 4.35 10^3^*µ*.

Despite this linear approximation, recognition is most probably not a constantly growing effect, and saturation must exist. However, following Rooney et al. data we understand that the tumor-immune interaction is beyond this region of reactivity saturation, so that we have build a saturation behavior to happen probably beyond *µ* ~ 10^−4^, a mutational level higher than those of most tumors measured by recent methodologies (see e.g. [24]).

## III. MATERIALS AND METHODS

### A. Population Dynamics of the tumor-immune interaction

The ecology of the cancer-immune system interaction pervades several complexity levels, from a vast antigenome [27] to multilayer cellular competition dynamics [28], and a first step towards modeling such ecology lies in dissecting which specific ingredients are key drivers in the phenomena we aim to understand.

Recent research points out that there might be up to 28 immune cell types with both antitumor and immuno-supressive roles infiltrated within a tumor [29]. Focusing on the immuno-surveillance mechanism of tumor growth inhibition following immune system recognition (early introduced in [30]), a minimal modelling approach recalls at least considering a population of tumor cells growing in competition with immune cells. This immune cohort represents a generic description for the global behavior of *cytotoxic immune cells*, such as CD8^+^ T cells (Fig. 1).

Despite further models have been useful at depicting very advanced properties of the immune system [31], we have chosen to keep a minimal scenario able to describe the competition dynamics at play. A complete description of the mathematical components of the model can be found in the Quick Guide to Model and major Assumptions toolbox. We use a well characterized model (see e.g. [32]) that has been used to describe clinical studies of cancer immunology such as tumor dormancy and tumor regression. This model has been studied using parameter ranges measured from experimental setups (Table I, see [20]).

### B. Ecological trade-offs in genetic instability

As discussed along the introduction, genetic instability plays a key role in tumor evolution, acting as the driving mechanism towards phenotypic variation and adaptation. Within our model, this can be translated as the replication rate being a function of its level of genetic instability *µ*. On the other hand we, *ρ*, the rate of cancer cell recognition by T cells, is also *µ*-dependent because of neoantigen production. Below we propose a minimal characterization of *r* and *ρ* able to describe how genetic instability modulates such trade-off.

#### 1. Cancer adaptation as a function of genetic instability

Cancer adaptation, here summarized to modulations in its replication rate, stems from the phenotypic plasticity resulting from mutations and copy-number alterations. On a general basis, enhanced tumor replication follows from mutations affecting oncogenic pathways, which poses a trade-off on genetic instability as it can, as well, damage any of the necessary machinery for cell viability.

Following previous research [22], an adaptive landscape is build on several assumptions based on the probabilities of mutating oncogenic and house-keeping genes (see Quick Guide to Model and Major assumptions). This adaptive landscape is of course of qualitative nature, and realistic fitness landscapes for unstable tumor environments are still far from our knowledge. However, certain points can be made if we give values within realistic parameter ranges to our function. The number of both oncogenes and house-keeping genes have been widely assessed, and we take them to be about *N_R_* ≈ 140 [33] and *N_HK_* ≈ 3804 [34] respectively. Interestingly enough, considering small replication effects for *δ_R_*, such experimental values produce an adaptive landscape that has an optimal region for tumor replication at about *µ* ≈ 10^−5^ − 10^−4^, which is in accordance with the point-mutation probability levels experimentally measured for unstable tumor cells [35].

#### 2. Immune recognition of malignancy as a function of genetic instability

Building a mathematical description of how the immune system reacts at the mutational burden of cancer cells is not straightforward. This stems from the fact that such behavior is yet starting to be understood at the molecular level and it probably builds upon many layers of complexity [8]. In our minimal mathematical approach, the first consideration is describing immune reactivity as proportional to recognition *ρ*, a rate that itself depends on the dynamics of neoantigen expression. Under our assumptions, since immune response follows from neoantigen detection we expect *ρ* being a function of the overall mutational landscape of a tumor, *µt*, which is eventually responsible for such neoantigen dynamics. Following recognition probability distributions from [19], we expect the average dominancy to initially increase with mutations as more and more neoantigens are generated and eventually saturate as very dominant neoantigens are rare.

The mathematical shape of this dependency *ρ*(*µt*) could stem from purely stochastic dynamics, but recent research gives better insight into the shape of this correlation. Rooney and colleagues provided an enlighting perspective in this direction by means of comparing a measure of immune response, following from the transcript levels of two key cytolytic effectors, with the total mutation count for eight tumor types [24].

Cytolytic response strengths in [24] seem to indicate a dependency on tissue and tumor microenviroment (Fig. 2a), which we have not included in our study since our model is not tumor type-specific. However, the shape of the immune response does seem to obey a global behavior across many cancer types, once cytolytic response values are normalized for each tumor type (Fig. 2b). With large outliers taken off from the data a global linear relation can be found for which normalized cytolytic activity scales with mutational load as CYT 4.35 10^−4^*µt* when averaged across the studied set of tumor types.

**FIG. 2:**
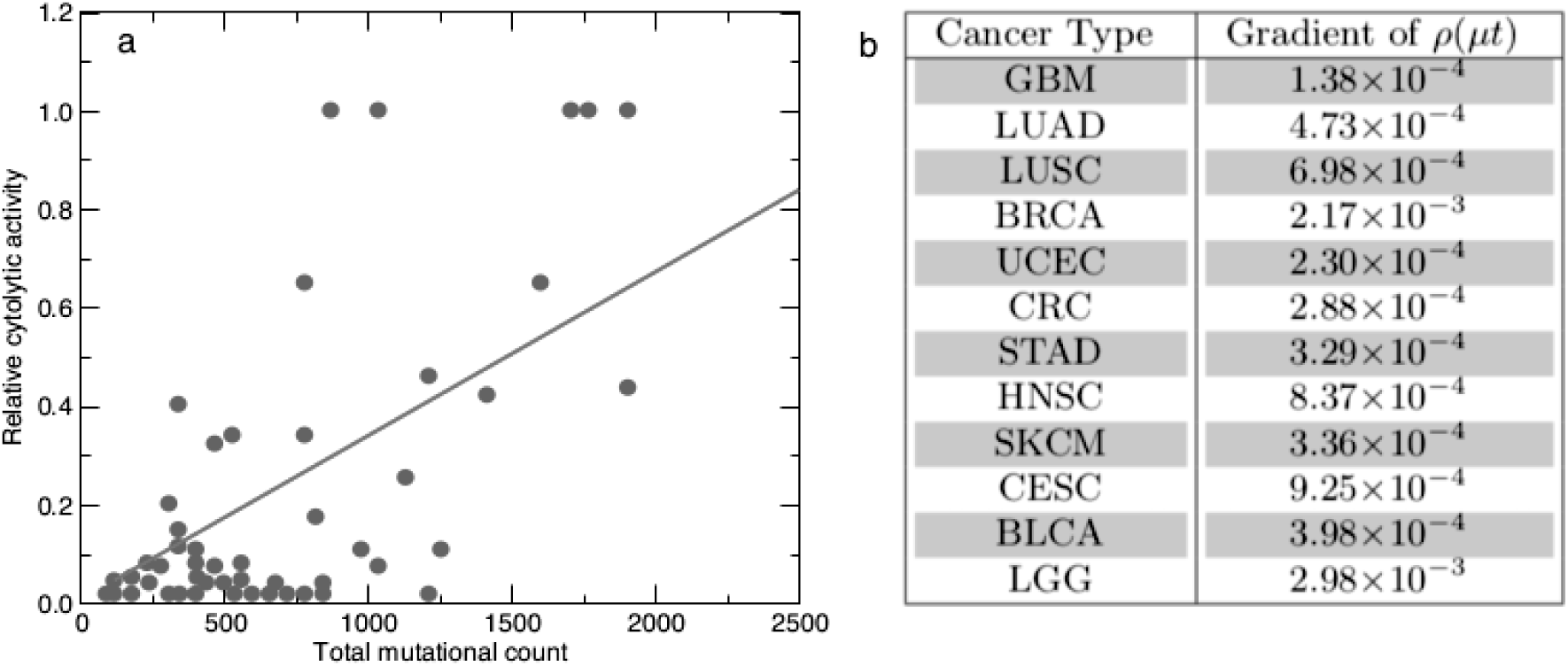
Measuring immune reactivity as a function of the mutational load. For 12 tumor types, a linear correlation between total mutation count and relative cytolytic activity is evaluated (Melanoma depicted in (a)). A linear function for *ρ*(*µt*) is obtained from averaging over all correlation strengths (b), resulting in *ρ*(*µt*) = 4.35 ⋅ 10^−4^*µt*. Data is directly obtained from [24]

Following assumptions on average tumor lifetime and cytolytic response saturation (see the Quick Guide to Model and Assumptions), we obtain a saturating function for *ρ*(*µ*) (with saturating curves not affecting the outcome of the model), and we can compare it with tumor adaptability *r*(*µt* (Figure 3) to obtain a full onedimensional landscape for tumor instability in the presence of T cells. Finally, let us mention as well that the model considers the cancer death rate, *δ*_c_, to be a linear function of *ρ*, since tumor cells will die out at rates depending on the strength of immune recognition.

**FIG. 3:**
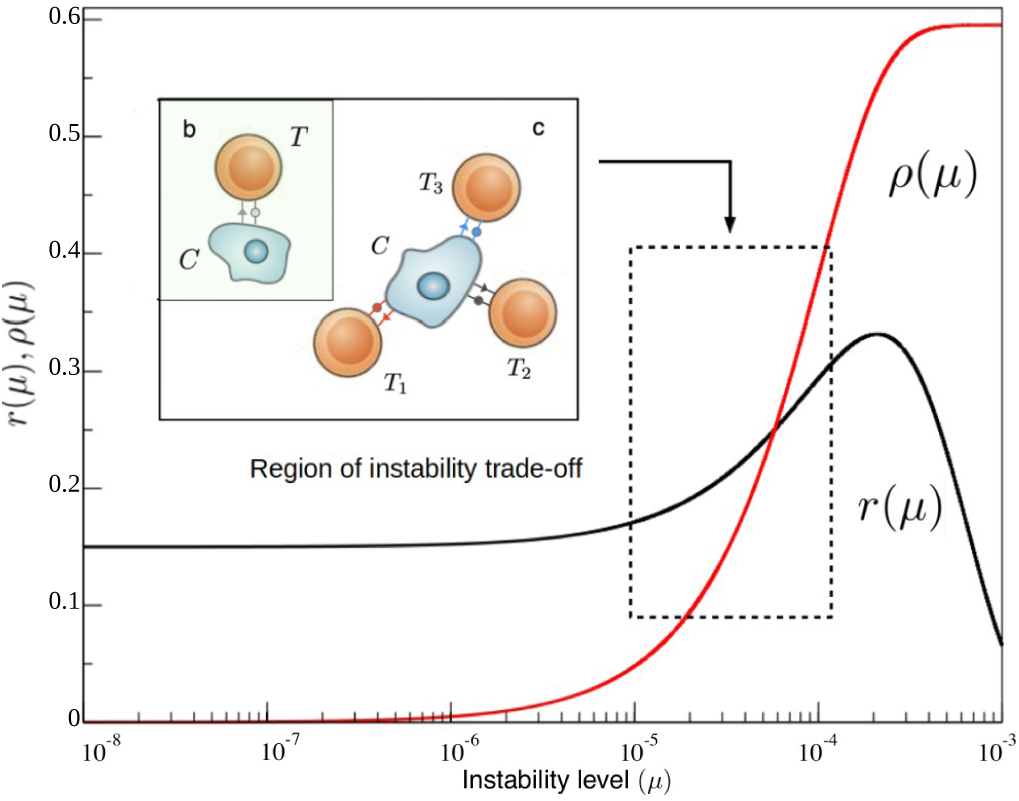
Functional forms for cancer replication *r*(*µ*) and immune recognition *ρ*(*µ*) functions. The first (black curve) provides a representation of the cancer instability landscape, as predicted from our theoretical approach (see Quick Guide to Model and Assumptions) and calibrated by available data. It reveals a very slow increase (in this log-linear diagram) at low instability levels followed by an increase associated to favourable mutations allowing for faster replication and a marked decay at high instabilities due to mutations on viability genes. The immune reactivity function *ρ*(*µ*) (in red, obtained from [24]) rises from zero to saturation beyond *µ* ~ 10^−4^. The relevant domain of common cancer instability levels is highlighted.

#### 3. Cancer-Immune system attractor states

Once discussed how to define the proper role of genetic instability on cancer adaptation and immune response, the original model is reinterpreted as a pair of coupled populations with instability-dependent rates, i.e.

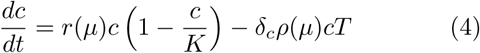

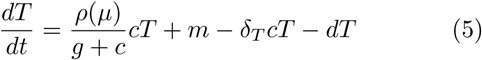

A global picture for the behavior of the system is obtained by studying its possible attractor states taking into account the variability of the mutational load. Together with the always present cancer free attractor (*c^∗^, T ^∗^*) = (0, *m/d*), other attractors can be inferred from studying the possible intersections between nullclines

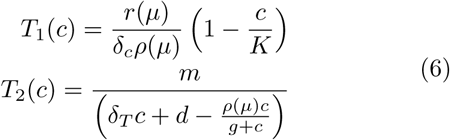

Nullcline 1 is a simple line with a negative slope controled by the inverse of the carrying capacity of cancer cells. On the other hand, nullcline 2 is a peaked curve, of height controled by immune cell migration and a denominator that might eventually produce divergences.

Through their crossings we will understand which steady states coexist under which parameter domains (See Results section and Figure 4).

**FIG. 4:**
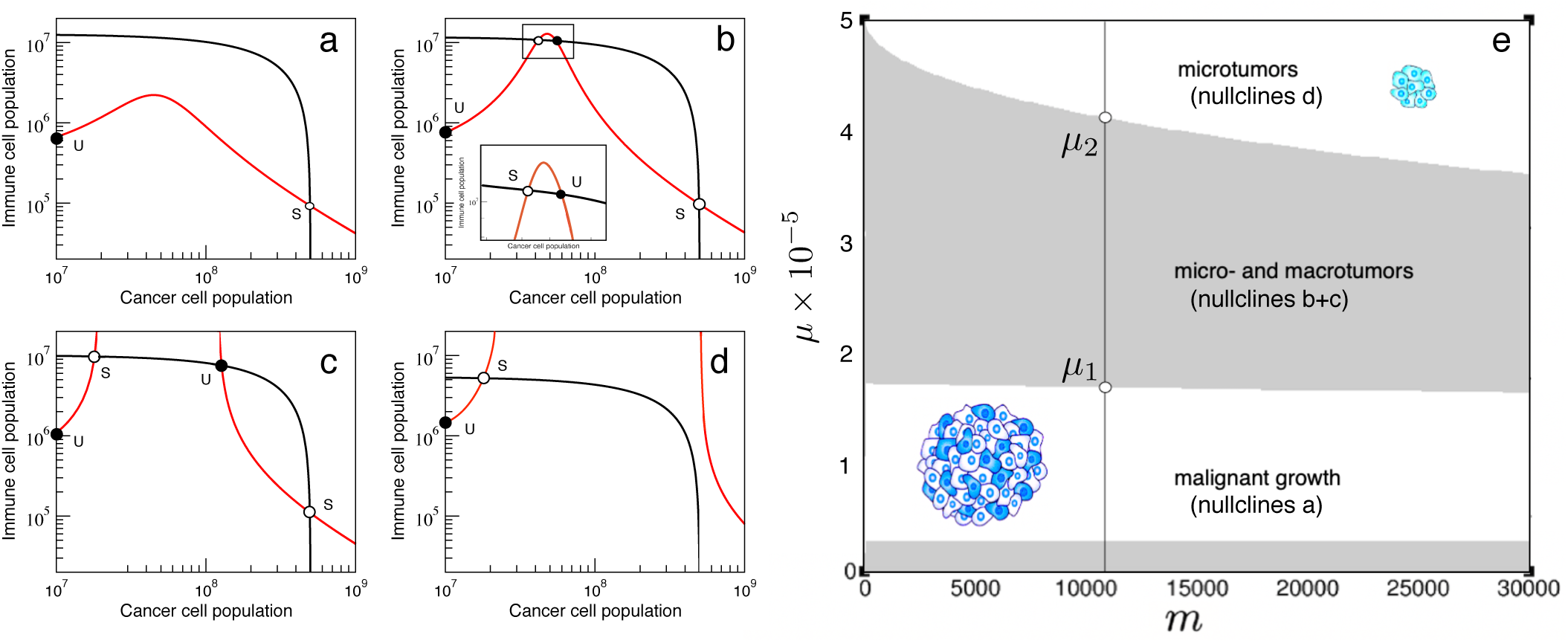
Cancer-Immune response attractors driven by instability. In (a-d) we display the nullclines as we increase mutation probability values. Two transitions can be seen. (a) At low genetic instability levels of 10^−5^ mutations per gene per division, such as those common in mutator tumors, only a large cancer attractor coexists with the unstable tumor-free equilibrium. (b) Beyond *µ^∗^* ~ 1.6 ⋅ 10^−5^, two new attractors are created, which correspond to a stable microtumor attractor and an unstable twin [32]. (c) At *µ^∗^* = 2.0 ⋅ 10^−5^, the microtumor attractor becomes smaller; until eventually the attractor of uncontrolled tumor growth is eliminated (d) at mutational levels similar to those attained after Mismatch-Repair knockout [36]. In (e) we summarise the bifurcation diagram for the possible scenarios as a function of *µ* and *m*.

Along with genetic instability, another parameter is key to the dynamics of the system. Regarding the second nullcline, we can see its size is linearly affected by the influx *m* of immune cells arriving at the tumor site. It is therefore interesting to understand how *µ* and *m* are related to cancer-immune scenarios, since this will open the door to further discussion on mutagenic and immune activation therapies.

By solving *T*_1_(*c*) = *T*_2_(*c*), we can understand how the values of *m* and *µ* affect the nature and number of possible solutions of the system. The previous identity leads to a cubic expression of the form *Ac*^3^ +*Bc*^2^ +*Cc* +*D* = 0, with

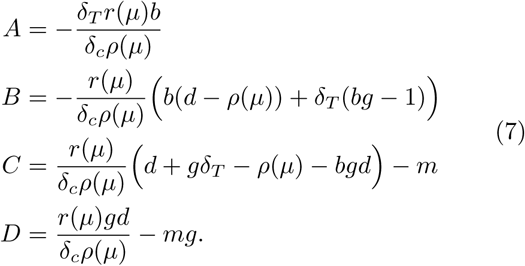

The sign of the discrimant Δ = 18*ABCD* – 4*B*^3^*D* + *B*^2^C^2^ – 4*AC*^3^ – 27A^2^D^2^ will give a notion of which binations of *m* and *µ* belong to which scenarios of figure 4. Knowing that three real roots exists for ∆ *>* 0 and only one for ∆ *<* 0, the transitions between attractor scenarios happen to occur at ∆ = 0. This condition can be used to easily describe the whole bifurcation space as seen in the results and figure 4e, giving a clear notion of how mutation frequencies and immune stimulation affect the possible outcomes of the system

## IV. RESULTS

### A. Minimal mutation rate for an efficient immune response

Before engaging into a full analysis of the complete model, we can study the behavior of the system for initial phases of progression. This corresponds to a small tumor of size *c* << *K* = 2 · 10^9^ cells. Under this assumption, the population dynamics of *c*(*t*) simplifies to

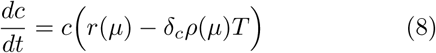

from where we can isolate a condition for tumor control, *i.e.*:

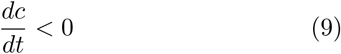

which leads to a crude estimate of the amount of T-cells required to counterbalance tumor growth

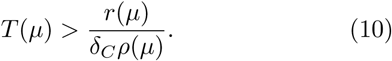

The inequality consistently shows that *T* (*µ*) will be proportional to the instability landscape of cancer growth rate divided by immune-mediated death. Using validated data from ([20], Table 1) and considering a healthy immune population of 10^7^ cells (Perelson and following sections), we obtain that the immune control condition is fulfilled for *µ* > 5.75 · 10^−5^ mutations per gene and replication. This preliminar result can be understood as a minimal mutation rate to generate a sufficient neoantigen load for immune attack. The estimated value is consistend within the range of genetic instability levels associated to MMR knockout [36].

**TABLE I:** Parameter values for the cancer-immune ecology model, estimated from experimental data of BCL_1_ lymphoma in the spleen of chimeric mice (see [20])).

| Parameter | Meaning | Kuznetsov et al (1994) estimate |
| --- | --- | --- |
| $r$ | Cancer cell replication rate | $0.18\text{day}^{-1}$ |
| $K$ | Tumor carrying capacity | $2 \cdot 10^9 \text{ cells}$ |
| $\delta_c$ | Immune-mediated cancer cell death rate | $1.101 \cdot 10^{-7} \text{day}^{-1} \text{cells}^{-1}$ |
| $\delta_T$ | Cancer-mediated immune cell death rate | $3.422 \cdot 10^{-10} \text{day}^{-1} \text{cells}^{-1}$ |
| $m$ | Rate of T cell migration towards tumor site | $1.3 \cdot 10^4 \text{ cells day}^{-1}$ |
| $g$ | Tumor size limit for effective T cell infiltration | $2.019 \cdot 10^7 \text{ cells}$ |
| $\rho$ | Rate of cancer cell recognition by the immune system | $0.1245 \text{day}^{-1}$ |
| $d$ | Intrinsic T cell death rate | $0.0412 \text{day}^{-1}$ |

### B. Transitions to tumor control and eradication at genetic instabilities within the MMR-knockout range

For well-formed tumors, no similar approach can be performed, but we can study the effects of changes in genetic instability in the sytem defined by equations (4) and (5) by picturing the intersections between nullclines described in the Materials and Methods section. It is straightforward to see how several transitions regarding creation and anihilation of steady states are governed by mutational probability *µ* (Fig. 4).

As expected from [32] and previous discussions, we know that the cancer-free attractor will always be present, but local stability will be ensured if *r*(*µ*)*/ρ*(*µ*) *< mδ_c_/d*, which only happens at very high instability levels of about 10^−3^ mutations per gene per division, and so no complete tumor clearence seems possible at realistic mutation rates for fixed *m*. Together with it, we can see that a large-tumor solution *c_L_* is also present at low instabilities (Fig 4a), and it is globally asymptotically stable. Interestingly enough, a transition seems to occur as the value for *µ* becomes larger: before *T*_2_(*c*) diverges, a smaller stable attractor *c_S_* is created together with its unstable *twin* (Fig. 4b), which is often described as a microtumor controlled by the immune system. Furthermore, nullcline 2 diverges at *µ* ~ 1.75 · 10^−5^ (Fig. 4c), and, as the two values for divergence of *T*_2_(*c*) grow further appart, the large cancer attractor disappears and only the controlled microtumor coexists with the cancer free attractor and is globally asymptotically stable (Fig. 4d). These results are consistent with those of [32], where such solution is considered a microtumor controled by the immune system. However, both transitions of microtumor creation and large tumor elimination being a function of the mutational levels of the tumor population are new to the present work.

At this point it is clear that understanding at what instability conditions these transitions happen is key to the possible outcomes of the tumor-immune interaction. For the given parameter region, a preliminar computational approach lets us see that the first transition happens around *µ* ~ 1.65 · 10^−5^ (Fig 4.b), whereas another transition where the large tumor attractor disappears happens at higher *µ* values of about *µ* ~ 4 · 10^−5^ (Fig 4.d).

Following extensive data, unleashed genetic instability after Mlh1 knockout in mice accounts for increasing mutation frequencies ranging from 10^−6^ ~ 10^−5^ up to 10^−4^ mutations per gene per division (values assessed for transgenic mice containing *supFG1* or *cII* from [36]). Interestingly enough, instability levels before MMR knockout put our system within a region where the large cancer attractor is stable and no controlled microtumor exists. However, the increase after Mlh1 knockout might be pushing cancer cells to a region beyond 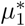, where the stable microtumor attractor appears, or even 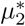, where the stable large cancer attractor has disappeared.

The resemblance between the model here discussed and experiments linking genetic instability to immune surveillance seems intuitive enough. Following [15], we think that there is a connection between the observed phenomenon of immune reactivity and tumor collapse after Mismatch Repair knockout and the qualitative behavior of our model, which depicts a transition of this kind at high *µ* values. Furthermore, we have taken advantage of recent research in order to use quantitative data to build our model. The fact that our model predicts the range for which immune surveillance reacts at increased cancer instability levels emphasizes the possible existence of transitions like the ones here studied.

Assessing if these two transitions are in fact well differentiated in vitro or if genetic instability can modulate tumor evolution towards controlled states can shed new light into the precise nature of mutagenic therapy as a mechanism towards increasing tumor immunogeneicity. Such therapies have produced key results in the field of virology [37], but within the context of cancer, drug design or resistance mechanisms have yet to be assessed, together with the problem of instability increases in non-mutator cells possibly generating secondary tumors [38].

#### C. Effects of modulating immune migration

Together with understanding the role of genetic instability, another key observation is considering *m*, the rate of immune migration, as a measure of immune activation. Further modeling, such as modulation and periodicity of treatment has already been studied [39], but studying the global scenario of how *µ* and *m* govern the cellular dynamics of the system might lead to further insight into possible combination therapies [7].

In the previous section we ensured the cancer free equilibrium would only be stable if *r*(*µ*)*/ρ*(*µ*) *< mδ_c_/d*, which is fulfilled at unattainable mutation rates of 10^−3^ mutations per gene and division. However, once we consider both *m* and *µ* clinical variables that can be affected by therapy, it is interesting to see that increasing *µ* produces an exponential-like decrease on the necessary migration rate needed to ensure the cancer-clearance scenario is stable.

Moreover, by picturing the bifurcation diagram as described in the MM section (Fig. 4), it is interesting to see how the first transition towards microtumor creation, 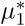, has a weak dependency on *m*, since the appearance of the intermediate attractors depends mostly on the denominator of nullcline 2 becoming null, so that *T*_2_(*c*) diverges at

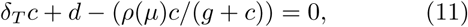

which is not a function of *m*. On the other hand, the transition to disappearance of the large-cancer attractor does depend on *m*, since *m* affects the width of *T*_2_(*c*), so that for higher *m* values *T*_2_(*c*) will go faster towards infinity and not cross *T*_1_(*c*). However, it seems intuitive from observing figure 4e that the role of genetic instability in increasing neoantigen production might be crucial even in the presence of high immune activation.

Mathematical work previous to our instability-driven model developed interesting considerations on derivation of cancer vaccines (see e.g. [40]), and introduced time dependent treatments [39] or time-delays in the immune response [41] based on the immune migration parameter, despite mathematical considerations remained somehow distant from clinical immunology and not many of the described behaviors after mathematically designed therapies have been observed *in vivo* [21].

In our model, it is first interesting to understand how, following parameter values taken from literature (Kuznetsov), the cancer-clearance scenario is hardly achievable by means of increasing only *µ* or *m*. However, therapy modulating both parameters would more easily bring the system closer to total eradication equilibria.

Together with this, it is seen that slight increases in immune migration *m* have a minor effect, compared to those of the genetic instability jump corresponding to MMR knockout, in producing other state transitions. However, recent advances in the field of adoptive cell transfer are resulting in the ability to grow up to 10^11^ antitumor lymphocites in vitro and select them for high-avidity recognition [42], which results in a whole novel scenario in the studied dynamics.

Recent research has highlighted the importance of genetic instability as a marker for good prognosis in immune checkpoint inhibition therapies [12,13]. Its role in neoantigen production is acknowledged as crucial [8]. Our results describing *µ* as another driver towards surveillance complementing *m* further support the relevance of genetic instability in the tumor-immune dynamics and complement the concept of cross therapies increasing tumor immunogeneicity [7].

## V. DISCUSSION

In the present work we have studied a minimal mathematical scenario describing how genetic instability, by means of enhancing tumor adaptation along with neoantigen production and immune recognition, can trigger sharp transitions towards tumor control and eradication.

Starting from basic considerations, we have asked ourselves about the ecological interactions between malignant cells and, in particular, effector immune cells able to respond after neoantigen recognition. Specifically, we consider how genetic instability, here as a mutation probability, shapes tumor adaptability and immune response.

Interestingly enough, genetic instability governs the possible outcomes of the system. Increasing mutational levels drives the system across two phase transitions. In the first one, two attractors are created consisting in smaller tumors coexisting with a larger population of T cells. This state has been characterized as a controled, but not totally eliminated microtumor [32]. The second transition accounts for the disappearence of the cancer-wins scenario, so that only solutions of immune control are present at large genetic instability levels.

Recent advances in the field of cancer immunology have proven that genetic instability is a key ingredient of the immune response [12–14], and particular research claims immune surveillance after MMR knockout follows from this causal relation between high mutational loads and neoepitope production [15]. Our model provides a conceptual description for how a transition between cancer growth and arrest can follow only from damaging DNA repair mechanisms.

Furthermore, we have used available data to calibrate the model parameters and to construct the immune recognition function. Using this information, we consistently explain phase transitions happening at microsatellite instability levels that resemble those of MMR knockout.

We have also studied the role of *m*, a parameter refering to immune migration or an eventual immune therapy. The model provides insights into how cross-therapies modulating both *m* and *µ* might be more effective in cancer clearance. We suggest that the relevance of *m* in producing the before mentioned transitions to tumor arrest is low, while minor increases in genetic instability seem much more effective at eradication of large tumors. It might be interesting to further discuss the possible meaning of this asymmetry between genetic instability and immune migration, as cross therapies inducing DNA damage prior to immunotherapy might proof much more effective [16]. We therefore postulate a possible mathematical description of recent discussions for novel perspectives on combination immunotherapy [7].

It is of course relevant, however, to understand how all the previous conclusions stem from a very minimal mathematical model, whereas the immune system is known to be a complex system of many interacting cell populations, whose role in cancer is not as straightforward as that of the particular effector T cells that we have here modeled [43,44].

Together with the fact that we have started from a simple two populations model, mainly because further models would generate hardly treatable systems [21], it is important to recall how *r* and *ρ* are functions of *µ* stemming from different approaches. Cancer adaptation is build on a simple probabilistic approach, discussed in previous research, while neoantigen recognition follows from a fit of disperse data together with an average over several tumor types and lifetime. This eliminates measures of inter- and intra-tumor heterogeneity. Despite considerable numerical approximations, the qualitative behavior stemming from our model is robust to changes in *r* and *ρ*.

Finally, as a result from the lack of heterogeneity, our model does not yet capture immune editing, a phenomenom at the core of immunotherapy failure, in which the tumor might develope immune resistance by means of either buffering the growth of immunosilent cells or editing its genome to express fewer neoantigens [45]. Using an evolutionary framework such as adaptive dynamics (cite aguade), future work might help to characterize in which regimes do cancer subclones evade immune surveillance through evolving their neoantigen landscape [46].

## Acknowledgements

The authors thank Elisa Beltran and Jordi Piñero for fruitful discussions, as well as the Complex Systems Lab members and Pierre Menard for their inspiring ideas. G.A. thanks Noemi Andor and the team at the *Evolutionary Biology and Ecology of Cancer Summer School 2018* for interesting comments on the cancer-immune system interaction. This work has been supported by the Botín Foundation by Banco Santander through its Santander Universities Global Division, a MINECO grant FIS2015-67616 fellowship, by the Universities and Research Secretariat of the Ministry of Business and Knowledge of the Generalitat de Catalunya and the European Social Fund and by the Santa Fe Institute.

## REFERENCES

1. Greaves M, Maley CC. Clonal evolution in cancer, Nature 2012;481:306–13

2. Lengauer C, Kinzler KW, Vogelstein B. Genetic instabilities in human cancers. Nature 1998;396:643–9.

3. Negrini S, Gorgoulis VG, Halazonetis TD. Genomic instability – an evolving hallmark of cancer. Nat Rev Mol Cell Biol 2010;11:220–8.

4. Andor N, Maley CC, Ji HP. Genomic instability in cancer: teetering on the limit of tolerance. Cancer Res 2017;77:1–7.

5. Sottoriva A, Spiteri I, Piccirillo SGM, Touloumis A, Collins VP, Marioni JC, et al. Intratumor heterogeneity in human glioblastoma reflects cancer evolutionary dynamics. Proc Natl Acad Sci U S A 2013;110:4009–14.

6. Miller Jf, Sadelain M. The journey from discoveries in fundamental immunology to cancer immunotherapy. Cancer cell 2015;27:439–49.

7. Zappasodi R, Merghoub T, Wolchok J. Emerging concepts for immune checkpoint blockade-based combination therapies. Cancer Cell 2018;33:581–98.

8. Schumacher TN, Schreiber RD. Neoantigens in cancer immunotherapy. Science 2015;348:69–74.

9. Efremova M, Finotello F, Rieder D, Trajanoski Z. Neoantigens generated by individual mutations and their role in cancer immunity and immunotherapy. Front Immunol 2017;8:1679.

10. Vesely MD, Schreiber RD. Cancer immunoediting: antigens, mechanisms, and implications to cancer immunotherapy. Ann NY Acad Sci 2013;128:1–5.

11. Segal NH, Parsons DW, Peggs KS, Velculescu V, Kinzler KW, Vogelstein B, et al. Epitope Landscape in breast and colorectal cancer. Cancer Res 2008;68: 889–92.

12. Rizvi NA, Hellmann MD, Snyder A, Kvistborg P, Makarov V, Havel JJ, et al. Mutational landscape determines sensitivity to PD-1 blockade in nonsmall cell lung cancer. Science 2015;348:124–8.

13. Snyder A, Makarov V, Merghoub T, Yuan J, Zaretsky JM, Desrichard A, et al. Genetic basis for clinical response to CTLA-4 blockade in melanoma. N Engl J Med 2014;371:2189–99.

14. Van Allen EM, Miao D, Schilling B, Shukla SA, Blank C, Zimmer L, et al. Genomic correlates of response to CTLA4 blockade in metastatic melanoma. Science 2015;aad0095.

15. Germano G, Lamba S, Rospo G, Barault L, Magr`ı A, Maione F, et al. Inactivation of DNA repair triggers neoantigen generation and impairs tumour growth. Nature 2017;552:116–20.

16. Le DT, Durham JN, Smith KN, Wang H, Bartlett BR, Aulakh LK, et al. Mismatch repair deficiency predicts response of solid tumors to PD-1 blockade. Science 2017;357:409–13.

17. Thorsson V, Gibbs DL, Brown SD, Wolf D, Borton DS, Ou Yang T et al. The immune landscape of cancer. Immunity 2018;48:812–30.

18. Bellomo N, Delitala M. From the mathematical kinetic, and stochastic game theory to modelling mutations, onset, progression and immune competition of cancer cells. Phys Life Rev 2008; 5:183–206.

19. Luksza M, Riaz N, Makarov V, Balachandran VP, Hellmann MD, Solovyov A, et al. A neoantigen fitness model predicts tumour response to checkpoint blockade immunotherapy. Nature 2017;551:517–20.

20. Kuznetsov Y, Makalkin I, Taylor M, Perelson A. Nonlinear dynamics of immunogenic tumors: parameter estimation and global bifurcation analysis. Bull Math Biol 1994;2:295–321.

21. Eftimie R, Bramson JL, Earn DJD. Interactions Between the Immune System and Cancer: A Brief Review of Non-spatial Mathematical Models. Bull Math Biol 2011;73:2–32.

22. Solé RV, Deisboeck TS. An error catastrophe in cancer? Jour Theor Biol 2004;228:47–54.

23. Aguadé-Gorgorió G, Solé RV. Adaptive dynamics of unstable cancer populations: The canonical equation. Evol Appl 2018;11:1283–92.

24. Rooney MS, Shukla SA, Wu CJ, Getz G, Hacohen N. Molecular and genetic properties of tumors associated with local immune cytolytic activity. Cell 2015;160:48–61.

25. Alexandrov LB, Nik-Zainal S, Wedge DC, Aparicio SAJR, Behjati S, Biankin AV, et al. Signatures of mutational processes in human cancer. Nature 2013;500:415–21.

26. Bozic I, Antal T, Ohtsuki H, Carter H, Kim D, Chen S, et al. Accumulation of driver and passenger mutations during tumor progression. Proc Natl Acad Sci U S A 2010;107:18545–50.

27. Charoentong P, Finotello F, Angelova M, Mayer C, Efremova M, Rieder D, et al. Pan-cancer immunogenic analyses reveal genotype-immunophenotype relationships and predictors of response to checkpoint blockade. Cell Rep 2017;18:248–62.

28. Hackl H, Charoentong P, Finotello F, Trajanoski Z. Computational genomics tools for dissecting tumour-immune cell interactions. Nat Rev Genet 2013;17:441–458.

29. Bindea G, Mlecnik B, Tosolini M, Kirilovsky A, Waldner M, Obenauf AC, et al. Spatiotemporal dynamics of intratumoral immune cells reveal the immune landscape in human cancer. Immunity 2013;39:782–795.

30. Ehrlich P. Collected papers in four volumes including a complete bibliography. Pergamon Press 1956.

31. Perelson AS, Weisbuch G. Immunology for physicists. Rev Mod Phys 1997;69:1219.

32. d’Onofrio A. A general framework for modeling tumor-immune system competition and immunotherapy: mathematical analysis and biomedical inferences. Physica D 2005;208: 220–235.

33. Vogelstein B, Papadopoulos N, Velculescu VE, Zhou S, Diaz LA Jr, Kinzler KW. Cancer genome landscapes. Science 2013;339:1546–58

34. Eisenberg E, Levanon EY. Human housekeeping genes, re-visited. Trends Genet 2013;29:569–74.

35. Tomlinson IPM, Novelli MR, Bodmer WF. The mutation rate and cancer. Proc Natl Acad Sci U S A 1996;93:14800–3.

36. Hegan DC, Narayanan L, Jirik FR, Edelmann W, Liskay RM, Glazer PM. Differing patterns of genetic instability in mice deficient in the mismatch repair genes Pms2, Mlh1, Msh2, Msh3 and Msh6. Carcinogenesis 2006;27:24022408.

37. Loeb LA, Essigman JM, Kazazi F, Zhang J, Rose KD, Mullins JI. Lethal mutagenesis of HIV with mutagenic nucleoside analogs. Proc Natl Acad Sci U S A 1999;96:1492–7.

38. Fox EJ, Loeb LA. Lethal mutagenesis: targeting the mutator phenotype in cancer. Semin Cancer Biol 2010;20:353–59.

39. Sotolongo-Costa O, Molina LM, Perez DR, Antoraz J, Reyes MC. Behavior of tumors under nonstationary therapy. Physica D 2003;178:242–53.

40. Gajewski T. Failure at the effector phase: immune barriers at the level of melanoma tumor microenvironment. Clin Cancer Res 2007;13:5256–61.

41. Galach M. Dynamics of the tumor-immune system competition: the effect of time delay. Int J Appl Math Comput Sci 2003;13:395–406.

42. Rosenberg SA, Restifo NP. Adoptive cell transfer as personalized immunotherapy for human cancer. Science 2015;348:62–8.

43. de Visser KE, Eichten A, Coussens LM. Paradoxical roles of the immune system during cancer development. Nat Rev Can 2006;6:24–37.

44. Chen DS, Mellman I. Elements of cancer immunity and the cancer-immune set point. Nature 2017;541:321–30.

45. Sharma P, Hu-Lieskovan S, Wargo JA, Ribas A. Primary, adaptive, and acquired resistance to cancer immunotherapy. Cell 2017;168:707–23.

46. Anagnostou V, Smith KN, Forde PM, Niknafs N, Bhattacharya R, White J, et al. Evolution of neoantigen landscape during immune checkpoint blockade in non-small cell lung cancer. Cancer Discov 2017;7:264–76.

